# Post-infection pig and ferret antisera show similar antigenic profiles for human influenza A(H1N1pdm09) viruses

**DOI:** 10.1101/2025.10.30.685573

**Authors:** Ruth Harvey, Tiphaine Cayol, Basudev Paudyal, Alice Lilley, Christine Carr, Catherine Hatton, Emily Briggs, Rodney S. Daniels, Samuel Richardson, Thomas P. Peacock, Nicola Lewis, Ian Brown, John W. McCauley, Elma Tchilian

## Abstract

**Background:** Monitoring antigenic drift in human influenza A viruses is essential for vaccine strain selection and ensuring protection against circulating strains. Antigenic drift is traditionally assessed using ferret antisera, which provide monospecific responses, and human vaccinee sera, which reflect exposure to multiple antigens. In this study we evaluated the pig as an alternative source of antisera to study antigenic drift compared to immune responses in ferrets and humans. We included seasonal influenza A(H1N1pdm09) human viruses that had shown different antigenic characteristics when using ferret or human antisera.

**Methods:** Pairs of pigs were inoculated with six human A(H1N1)pdm09 viruses circulating between 2019 and 2023, a period of marked antigenic drift. Pig and ferret antisera were analysed by hemagglutination inhibition (HI) and virus neutralization (VN) assays.

**Results:** Pigs were successfully infected with all strains, shedding virus and producing antibody responses, confirming their susceptibility to human influenza A viruses. Antigenic reactivity of pig antisera was qualitatively comparable to ferret antisera in both HI and VN assays, although maximum homologous antibody titres were significantly higher in ferrets. The antisera raised against viruses in circulation in 2019 and before, exempified by A/Guangdong-Maonan/SWL1536/2019, clade 5a.1, were clearly differentiated by both ferret and pig antisera from those in clade 5a.2 and its derivatives that became predominant.

**Conclusions:** Ferrets and pigs showed comparable responses and both distinguished clade 5a.1 from clade 5a.2. However, neither model recognised antigenically drifted variants from 2019–2022, including subclades 5a.2-C, 5a.2a-C.1/C.1.9, and .5a.2a.1-C.1.1/D, which were distinguishable using human post-vaccination antisera.

## Introduction

Human seasonal Influenza A viruses continually evolve genetically and antigenically necessitating constant monitoring and frequent updating of seasonal influenza vaccine strains. The Global Influenza Surveillance and Response System (GISRS), coordinated through the World Health Organisation (WHO), monitors the genetic and antigenic relatedness of currently circulating viruses, and each year the WHO makes separate recommendations of virus components for inclusion in seasonal influenza vaccines for both the Southern and Northern Hemisphere (SH and NH) [1].

Underpinning the vaccine strain recommendations are phenotypic data characterising the antigenic properties of currently circulating influenza strains relative to previously recommended Candidate Vaccine Viruses (CVVs) and to other reference viruses, including viruses chosen to be representative of emerging genetic groups. Routinely the ferret is used as a small animal model for antibody responses to human seasonal influenza and reference antisera raised against CVVs and other key reference viruses and used throughout each influenza season in hemagglutination inhibition (HI) and virus neutralisation (VN) assays to assess the antigenic properties of contemporary influenza virus strains. These ferret antisera antigenic analyses are supplemented by human serological response analyses using pre- and post-vaccination serum panels.

Periodically disparities have been observed between the relative ability of ferret and human immunological responses to similarly discriminate amino acid substitutions in hemagglutinin (HA) resulting in antigenic evolution, particularly with the A(H1N1pdm09) human influenza viruses. For example, while the ferret antibody responses to 2019 CVVs aligned with the human responses when assessed by HI assays, there were discrepancies with CVVs from 2021 such as A/Sydney/5/2021 (Syd21), where human serology indicated significant antigenic drift commensurate with a need to update the vaccine strain whereas ferret antisera did not [2, 3].

Pigs are an important host for influenza A viruses and are readily infected by the same H1N1 and H3N2 virus subtypes as humans with periodic transmission in both directions [4]. They also serve as a significant biomedical model for studying human disease [5]. Pigs have a longer life span, and are genetically, immunologically, physiologically, and anatomically more like humans than small laboratory animals [6, 7]. Additionally, pigs exhibit a comparable distribution of sialic acid receptors in the respiratory tract to humans [8]. Indeed we have previously shown that pig immune responses following infection with A(H1N1pdm09) influenza virus were similar to those induced in humans [9] thereby providing a model of human influenza infection [10].

Here we tested whether the pig immunological response was able to discriminate between human A(H1N1pdm09) influenza viruses that might be distinguished by either post-infection ferret antisera or post-vaccination human sera.

## Results

### Virus infection of pigs and nasal shedding

The panel of study viruses selected were representative of viruses in circulation between 2019 and 2023 (**Table 1**). Over this period some groups of viruses were differentiated by both post-infection ferret antisera and post-vaccination human sera while others had previously shown detectable differences in response in human serology studies but not when tested against post-infection ferret antisera. Six groups of two pigs each were infected intranasally with one of the study viruses possessing the HA amino acid substitutions indicated (**Table 1**) and sampled as described in Materials and Methods and illustrated in **Fig 1A**. All animals were productively infected as determined by nasal shedding of virus. The infection kinetics showed that mean shedding following infection with all viruses generally peaked at 4-days post-infection (DPI) (Syd21 infected pigs peaked at 2-DPI) and had been cleared in all pigs by 7-DPI (**Fig. 1B**). Virus whole genome sequencing of the input and 4-DPI nasal swabs showed all shed viruses to be unchanged at the consensus level from the infecting viruses, indicating lack of genetic mutations associated with pig infection in a single animal infection cycle (data not shown).

**Table 1.**
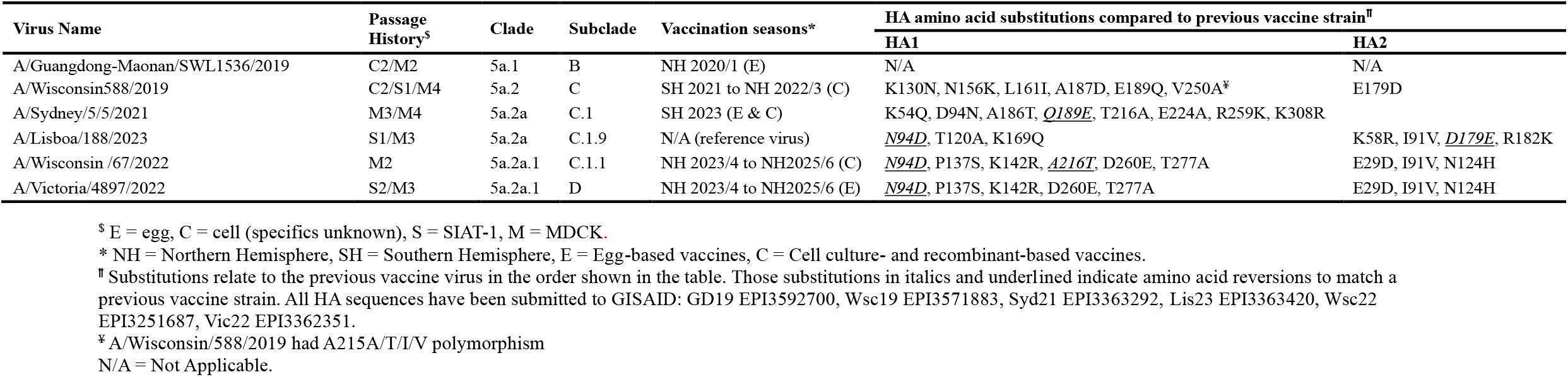
Influenza A(H1N1pdm09) viruses used in this study.

**Figure 1.**
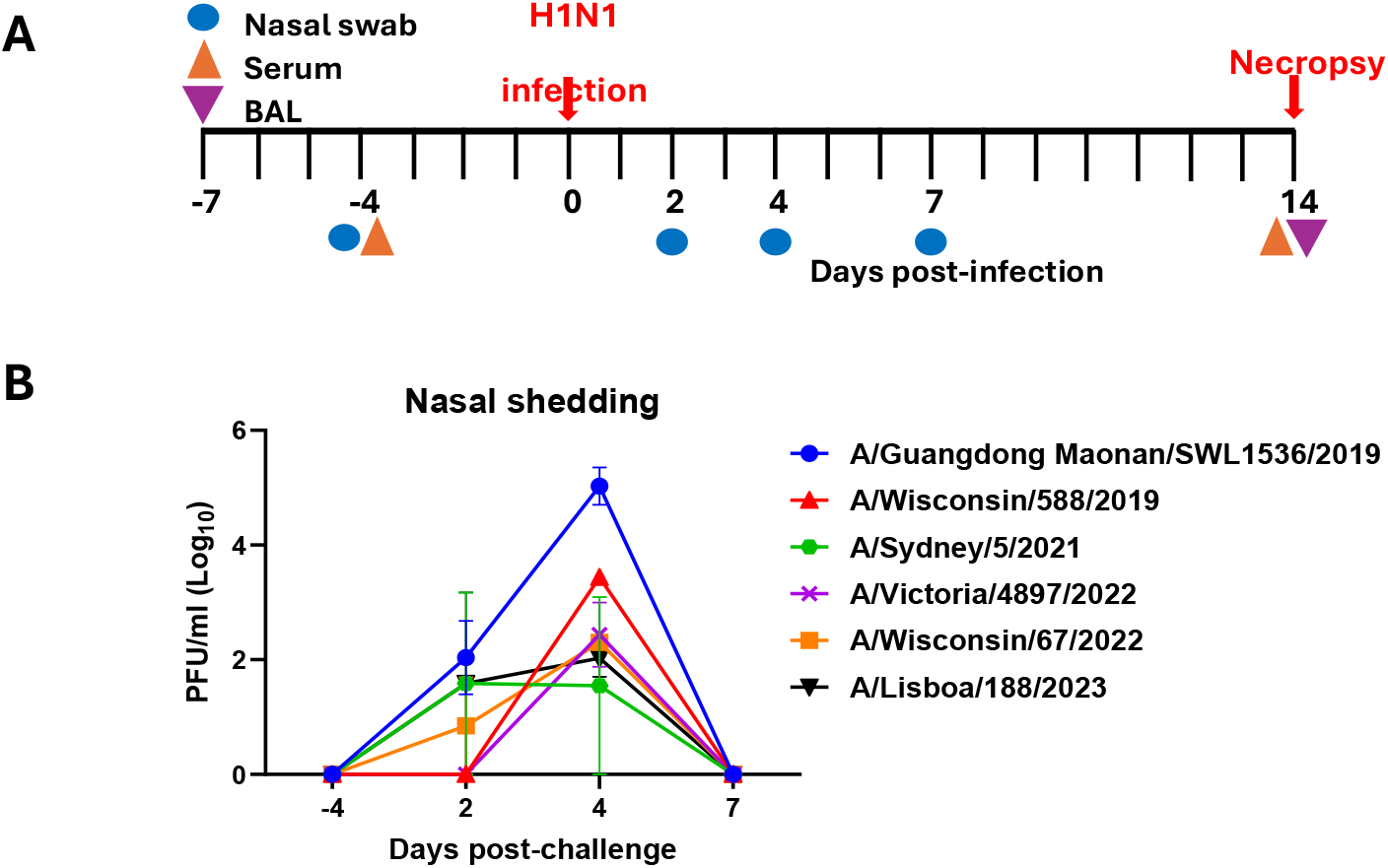
Experimental pig infection and analyses of samples taken. (**A**) Schematic of the experimental design is shown, two pigs each were intranasally infected with one of the A(H1N1pdm09) viruses shown in **Table 1**. Serum and nasal swabs were collected at indicated times with necropsy at 14-DPI. (**B**) Virus loads in nasal swabs at -4-, 2, 4- and 7-DPI were determined by plaque assay. Each data point represents the mean of the two pigs inoculated with the same virus and error bars show the standard deviation of that mean.

### HI responses of ferret and pig antisera

Antibody titres in HI showed differences in the magnitude of responses between ferret and pig antisera against the panel of study viruses (**Table 2A**). Ferret antisera consistently exhibited high titres of 5120 against the homologous viruses used for infection, indicating a strong antigen-specific response. In contrast, pig antisera generally demonstrated lower homologous titres, ranging from 160 to 640, depending on the specific virus and individual animal.

**Table 2.**
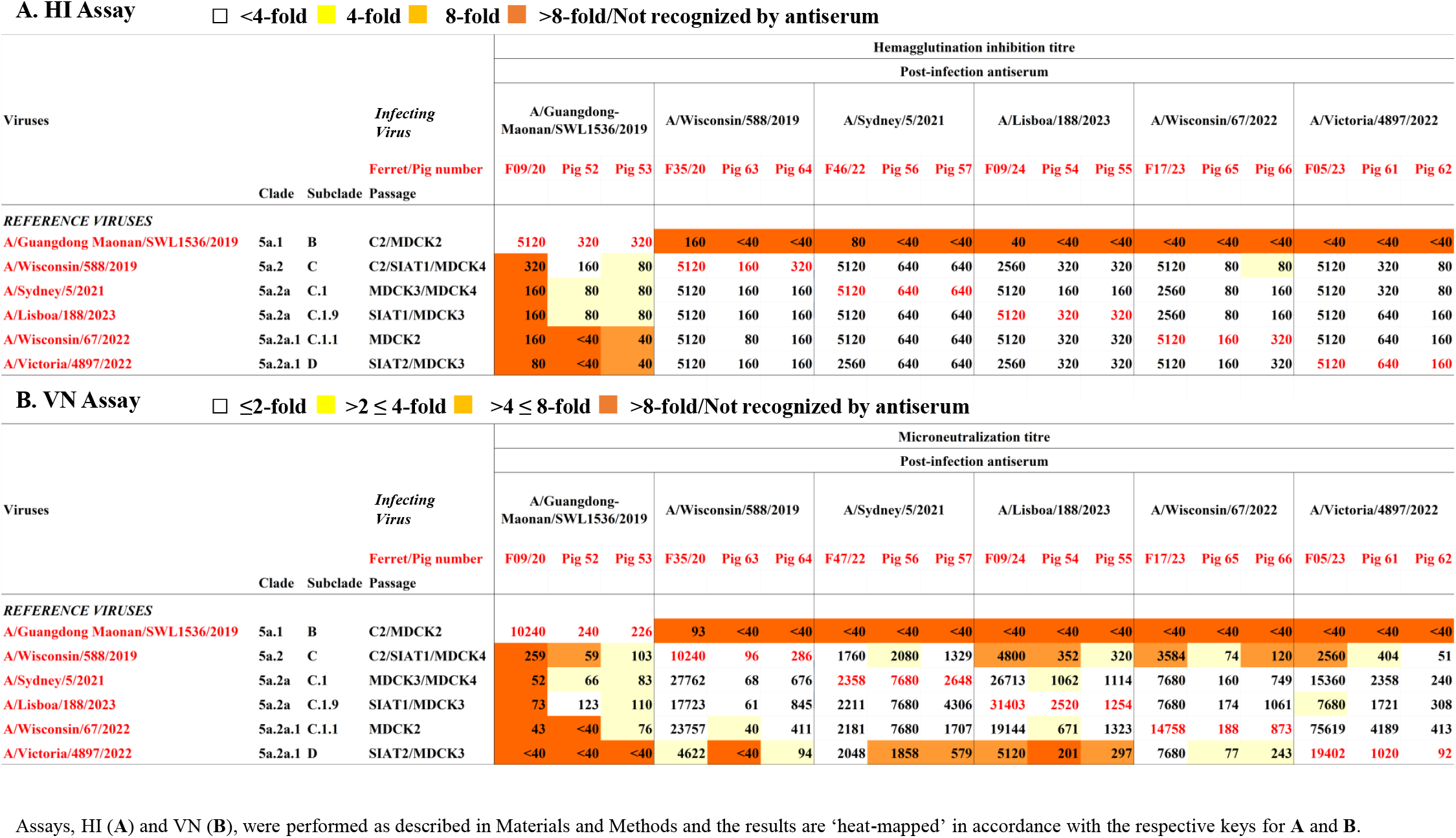
Virus neutralizing activity of ferret and pig antisera assessed by HI and VN.

Cross-reactivity patterns were broadly similar between ferret and pig antisera. The ferret antiserum raised against A/Guangdong-Maonan/SWL1536/2019 (GM19;subclade 5a.1-B) showed good recognition of the homologous antigen but very limited recognition of viruses of subclades 5a.2-C A/Wisconsin588/2019 (Wsc19), 5A.2a-C.1 (Syd21), 5A.2a-C.1.9 A/Lisboa/188/2023 (Lis23), 5a.2a.1-D A/Victoria/4897/2022 (Vic22) and 5a.2a.1-C.1.1 A/Wisconsin/67/2022 (Wsc22). Correspondingly, the antisera raised against Wsc19, Syd21, Lis23, Vic22 and Wsc22 all showed high HI titres for the homologous viruses and poor recognition of GM19. However, each of these antisera recognised viruses in subclades 5a.2-C, 5a.2a-C.1, 5a.2a-C.1.9, 5a.2a.1-D and 5a.2a.1-C.1.1 at titres within 2-fold of the homologous titres, most at titres equal to the homologous titre.

While there were some differences in the magnitude of HI antibody titres between individual pigs using the same infecting virus, two-fold differences were not considered significant since the antigenic profiles were consistent when comparing serum reactivity between homologous and heterologous viruses for each pair of pigs. One exception was a 4-fold variation in the homologous titres for Vic22 infected pigs (pigs 61 & 62) but titres to heterologous viruses showed identical reactivity profiles (**Table 2A**). Like the ferret antiserum raised against GM19, the two pig antisera raised against GM19 recognised the homologous virus well, at HI titres of 320, but recognised the heterologous viruses less well, although pig 53 and 54 antisera recognised Wsc19, Syd21 and Lis22 at titres within 4-fold lower than the homologous titres. In contrast, all antisera raised against the other five study viruses recognised GM19 poorly, but showed good recognition of the homologous and heterologous viruses, with a single exception for pig 66 (infected with Wsc22) antiserum which recognised Wsc19 at a titre 4-fold reduced compared to the homologous titre.

### VN of ferret and pig antisera

Ferret antisera raised against GM19 showed a similar clear differentiation in VN assays between the homologous virus and the other five study viruses, as seen by HI assay (**Table 2B**). Moreover, the antisera raised against the other five study viruses all recognised GM19 poorly. Nevertheless, these antisera showed more variable recognition of the heterologous viruses: those raised against Lis23 (5a.2a-C.1.9), Wsc22 (5a.2a.1-C.1.1) and Vic22 (5a.2a.1-D) all recognised Wsc19 (5a.2-C) at titres >4-fold reduced compared to homologous titres, as was the case for Lis23 antiserum with Vic22. Reductions of 2-4-fold were seen for Wis19 antiserum with Vic22 and Vic22 antiserum with Lis23.

Within pig antisera pairs homologous VN titres ranged from equivalence, for GM19, up to 5-fold (Wsc22) for four pairs while for Vic22 (pigs 61 and 62) 21-fold difference was seen (**Table 2B**). Neutralization titres were consistently higher in ferret antisera for all viruses except Syd21 which had homologous titres of 2358 in ferret and 7680 or 2648 in pigs. Further, the pig antisera showed an even greater variation in recognition patterns in the VN assay than was seen in the HI assay. However, heterologous titres were inconsistent between pig pairs in their profile of reactivity when directly comparing to homologous titres for all infecting viruses.

Again, pig antisera differentiated GM19 (clade 5a.1) from the five viruses in clade 5a.2 and derivatives thereof, antisera raised against them recognising GM19 poorly. Some neutralization of the heterologous viruses was seen by the antisera raised against GM19, five of the ten heterologous titres being within 4-fold of the homologous titres, one within 2-fold. Of the 40 heterologous titres associated with antisera raised against the other five study viruses, seven were >4-fold reduced: five related to Vic22 with antisera raised against Wsc19 (pig 63, with a low homologous titre), Syd21 (both pigs) and Lis23 (both pigs), and two to Wsc19 for antisera raised against Lis23 (pig 54) and Wsc22 (pig 66). Notably, Wsc19 was also recognized poorly by ferret antisera raised against Lis23 and Wsc22.

Overall, antisera raised in both pigs and ferrets against viruses in clades 5a.2, 5a.2a and 5a.2a.1 recognised the 5a.1 virus, GM19, poorly in both HI and VN assays. The antiserum raised against GM19 in ferrets recognised the 5a.2, 5a.2a and 5a.2a.1 viruses poorly in both assays, but the antisera raised in pigs recognised some of the viruses from the heterologous subclades at variable titres, many at titres within 4-to 8-fold of the titres for the homologous antigen. Antisera in ferrets and pigs raised against 5a.2, 5a.2a and 5a.2a.1 viruses recognised viruses in the same clades well, all at titres within 4-fold of the homologous titres, all but one within 2-fold, in HI assays. There was more variation seen in the VN assay but there was no observed differentiation between antigens in individual clades 5a.2, 5a.2a and 5a.2a.1.

### Human Serology

Three serum panels taken from adult volunteers (18 to 64 years of age) over three consecutive vaccination campaigns were analysed: 2020 ahead of the northern hemisphere (NH) 2020/21 season, 2021 for the NH 2021/22 season, and 2022 for the NH 2022/23 season. Volunteers were immunised with inactivated vaccine antigens corresponding to egg-propagated GM19 (clade .5a.1) in 2020, and cell culture-propagated Wsc19 (clade 5a.2) in 2021 and 2022. These antisera were used in HI assays with the homologous vaccine viruses and cell culture-propagated Wsc19 for the 2020 panel, and cell culture-propagated cultivars of Syd21 (subclade 5a.2a-C.1) and Wsc22 (subclade 5a.2a.1-C.1.1) for the 2021 and 2022 panels.

Non-parametric analysis of the paired post-vaccination HI titres using a Wilcoxon matched-pairs signed rank test showed that the human sera differentiated GM19 from Wsc19 (P< 0.0001) in the 2020 vaccinees. The sera from the 2021 vaccinees clearly differentiated Wsc22 from Wsc19 (P=0.0005) but not Syd21 (P=0.083), whilst the sera from the 2022 vaccinees differentiated both Syd21 (P=0.0009) and Wsc22 (P=0.0001) from Wsc19 (**Fig 2**).

**Figure 2.**
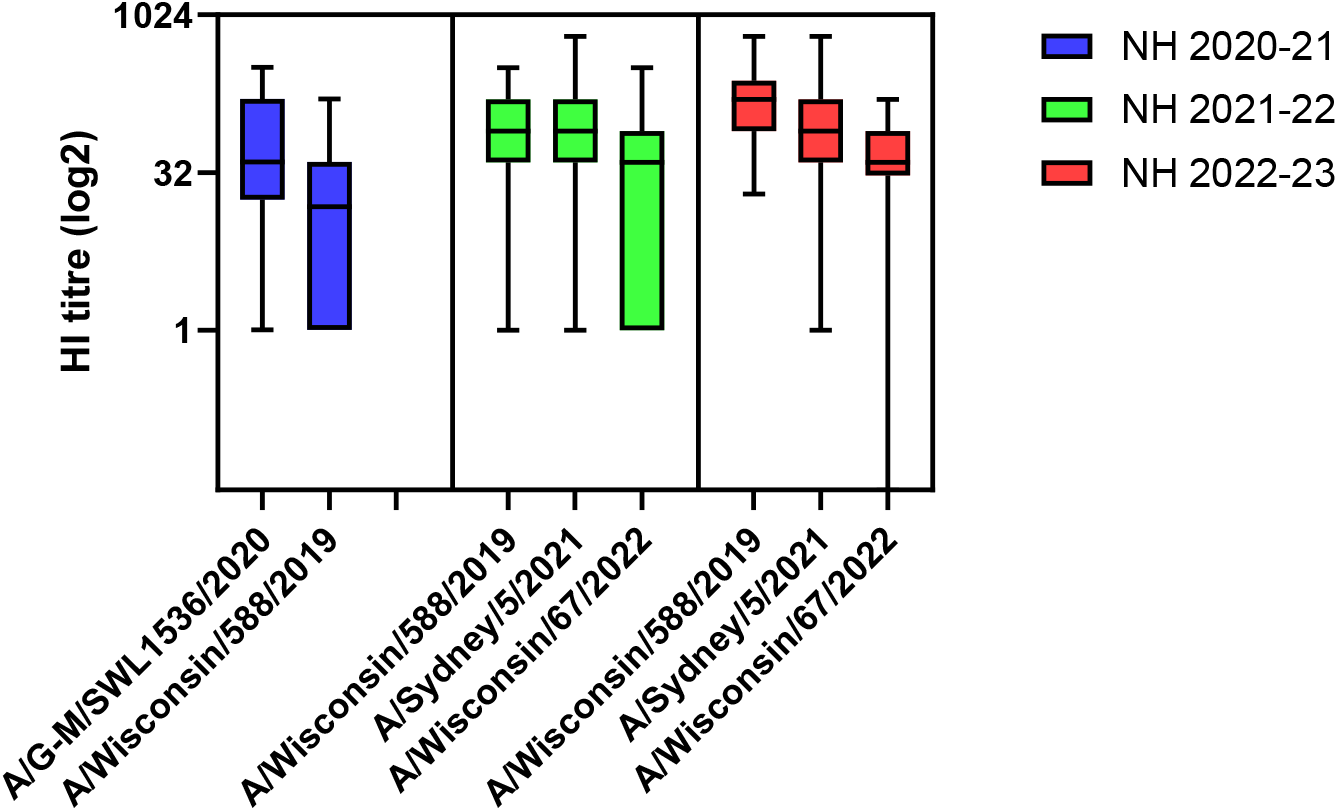
Post-vaccination HI titres of three serum panels from healthy adults. Serum panels from three consecutive northern hemisphere (NH) vaccination campaigns: 2020 for 2020/21 season (in blue), 2021 for 2021/22 season (in green), and 2022 for 2022/23 season (in red). Each panel was tested against the A(H1N1pdm09) virus component of the vaccine for the season and test viruses as shown. Data is plotted on a log_2_ scale.

Analysing individual serum responses showed that of the 22 volunteers vaccinated with egg-propagated GM19 eight recognised Wsc19 at titres reduced 4-fold or more. Of the 29 volunteers vaccinated with cell culture-propagated Wsc19 in 2021, 12 (41%) recognised Syd21 and 22 (76%) recognised Wsc22 at titres reduced 4-fold or more while of the 25 volunteers vaccinated in 2022, 9 (36%) recognised Syd21and 13 (45%) recognised Wsc22 at titres reduced 4-fold or more.

These results indicate that, like post-infection ferret and swine antisera, post-vaccination human sera (although with some degree of variation) are able to differentiate the clade 5a.1 vaccine virus from Wsc19 (clade 5a.2). Post-vaccination serum panels as a whole, also differentiate the vaccine virus Wsc19 from Wisc22 (subclade 5a.2a.1-C.1.1) and in one of the two seasons differentiated Wsc19 from Syd21 (subclade 5a.2a-C.1) unlike the ferret and pig antisera. These results with human sera are consistent with published results from post-vaccination human serum panels used in vaccine strain recommendations for the Southern Hemisphere 2023 and the Northern Hemisphere 2023/24 influenza seasons [2, 3]. Nevertheless, individual sera differed in their discrimination between viruses in the different subclades.

## Discussion

Post-infection ferret antisera and post-vaccination human sera are usually used to define the antigenic characteristics of human influenza viruses as they evolve. However, for some recent seasonal influenza viruses there has been a disparity in the recognition of viruses between human and ferret antisera which has impacted on human vaccine strain selection, notably so for A(H1N1pdm09) viruses [2, 11–13]. Therefore, this study aimed to evaluate the immune response in pigs following infection with A(H1N1pdm09) human influenza viruses, to compare their recognition of recent A(H1N1pdm09) viruses with those of ferret and human antisera, and to investigate whether post-infection pig antisera can enhance the antigenic analysis of human influenza A(H1N1pdm09) viruses.

The overall study findings revealed that pigs could readily be infected with seasonal human A(H1N1pdm09) influenza viruses. This was not unexpected and confirmed that pigs remain susceptible to contemporary human seasonal viruses descended from the 2009 swine origin pandemic virus and support ongoing detections of virus spillover from humans to pigs [14, 15]. Broadly, pig immune responses produced consistent antigenic profiles directly comparable to those of ferrets; although, we obtained some altered characteristics when comparing neutralization profiles for ferret and pig antisera between homologous and heterologous viruses, but these were not correlated with subclade.

The post-infection antisera from ferrets and pigs and the post-vaccination human sera distinguished GM19 (clade 5a.1) from Wsc19 (clade 5a.2). The HA of this pair had the substitutions K130N, N156K, L161I, A187D, E189Q and V250A in HA1 and E179D in HA2 (**Table 1**), and four of these seven substitutions can be mapped to antigenic sites: Sa (K130N and N156K) and Sb (A187D and E189Q). Five of these HA1 substitutions (not E189Q) were conserved in the remaining four members of the virus panel. The antisera from both ferrets and pigs did not differentiate Wsc19 from Syd21 (subclade 5a.2a-C.1) in HI or VN assays, with Syd21 having HA1 substitutions K54Q, D94N, A186T, Q189E (reversion), T216A, E224A, R259K and K308R (**Table 1**). These antisera also did not distinguish Vic22 (subclade 5a.2a.1-D), Lis23 (subclade 5a.2a-C.1.9) and Wsc22 (subclade 5a.2a.1-C.1.1) from Wsc19, despite viruses in these subclades differing in HA1 amino acid sequence at several sites including the substitutions T120A, P137S, K142R, K169Q, D260E and T277A (**Table 1**). Importantly antisera from both animal species did not clearly discriminate between Wsc19 and Wsc22 in contrast to the post-vaccination cohorts of human sera in 2021 and 2022, which showed a reduction in human sera reacting to Wsc22 (**Fig 2** and [2, 3]). This pair of viruses differed at nine HA1 residues K54Q, P137S, K142R, A186T, Q189E, E224A, R259K, D260E and T277A. The altered recognition by the post-vaccination sera resulted in a recommendation to update the cell-based and egg-propagated vaccine strain components for the NH 2023/24 season [3].

Although the pattern of recognition of ferret and pig antisera were similar, direct comparisons of the immune responses showed that while the pairs of pigs in this study had comparable homologous and heterologous HI titres they were lower than those of ferret (approximately 16-fold across all viruses, except for GM19, a virus antigenically very distinct from the other viruses examined) which reduces the dynamic range of the assays. When homologous titres in VN assays are compared, generally pig antisera titres were lower than those of ferret, showing pig-specific reductions of 12-to 211-fold, for all but antisera raised against Syd21 (**Table 2B**).

The utility of pig and ferret antisera in evaluating responses to phenotypic evolution of swine influenza viruses has recently been compared [17]. In that study swine sera were produced following intranasal infection and subsequent intramuscular immunisation with the same virus dose formulated with adjuvant, whereas ferret sera were obtained after intranasal infection only. Despite these differences in immunisation/infection regimes, the breadth and relative magnitude of the antibody responses were equivalent in both hosts, as assessed by HI and VN.

We have previously demonstrated that monoclonal antibodies (mAbs) derived from A(H1N1pdm09)-infected pigs recognized the same two major immunodominant HA sites, Sa (residues 160 and 163) and Ca (residue130) [9]. In 2014, A(H1N1pdm09) viruses acquired antigenic drift through HA1 K163Q substitution which was not detected using ferret antisera [11, 18–20]. Whether mAbs generated from the infected pigs in this study would reveal the fine specificity of epitopes in Wsc19 and Wsc22 remains to be determined. By 2016, the dominant A(H1N1pdm09) viruses circulating had acquired a NQS glycosylation motif through HA1 S162N substitution, effectively masking the Sa antigenic site.

These results reaffirm the continued importance of the interface between human and pigs for ongoing influenza virus evolution and potential antigenic divergence whilst retaining interspecies transmissibility. The genetic data obtained from viruses prior to infection and those shed four days after infection showed retention of consistent human-derived virus genetic characteristics, supporting maintenance of similar viruses in pigs albeit without defining transmissibility between pigs. Given the high susceptibility of pigs to human A(H3N2) influenza viruses and the maintenance of such viruses many years after the antigenic variants have disappeared from humans [21], due to genetic drift, there is also merit in assessing comparative data between ferrets and pigs to discriminate antigenic drift leading to A(H3N2) vaccine strain changes.

This is the first study directly comparing ferret and pigs as a host to define antigenic reactivity and variation in human A(H1N1pdm09) influenza viruses. We were able to show little variation qualitatively between the two animal hosts when using post-infection antisera to discriminate between viruses. Therefore, pigs could offer a qualified alternative for measuring antigenic variation in human influenza A viruses. This study also broadens the scope of pigs to serve as an animal model for human influenza for evaluating immune responses and vaccine studies as one of the milestones for improved influenza vaccine research [10].

## Materials and Methods

### Virus propagation

Viruses provided by the Francis Crick Institute (Table 1) were propagated in Madin Darby Canine Kidney (MDCK) cells. MDCK were cultured in T75 flasks with Dulbecco’s Modified Eagle’s Medium (DMEM; Sigma Aldrich), 10% fetal bovine serum (FBS; Gibco) and 1% Penicillin and Streptomycin (PS, Sigma Aldrich). Cells were inoculated with 2 ml of a seed virus diluted in DMEM+PS and incubated for 30 minutes at room temperature before adding 10 ml of DMEM+PS+2µg/ml (TPCK)-treated trypsin and incubated for two to three days at 37°C with 5% CO_2_. The gene sequence of each virus stock was confirmed before use.

### Influenza infection of pigs

Twelve 6-8 weeks old, outbred Landrace x Large White pigs (6 female and 6 male) were obtained from a commercial high-health status herd. The pigs were screened by ELISA for the absence of serum IgG titres against A(H1N1pdm09) and A(H3N2) virus. For each study virus two pigs, with one male and one female randomly divided per group, were infected. After a week of acclimatization the two pigs in each study group were inoculated intranasally using a mucosal atomization device (MAD, Medtree, UK) with 2 ml (1 ml per nostril) with either A/Guangdong Maonan/SWL1536/2019 (GM19; 1.8 × 10^5^ PFU/pig) or A/Lisboa/188/2023 (Lis23; 5 × 10^6^ PFU/pig) or A/Sydney/5/2021 (Syd21; 8 × 10^6^ PFU/pig) or A/Victoria/4897/2022 (Vic22; 4 × 10^6^ PFU/pig) or A/Wisconsin/588/2019 (Wsc19; 1 × 10^6^ PFU/pig) or A/Wisconsin/67/2022 (Wsc22; 1.5 × 10^6^ PFU/pig) (**Fig. 1A**). At 14 days post-infection (DPI), pigs were humanely euthanized with an overdose of pentobarbital sodium anaesthetic and clotted blood and bronchoalveolar lavage fluid (BAL) was collected to assess immune responses to the viruses.

### Sampling of study pigs

Nasal swabs (NS) (two per nostril) were taken from pigs four days before infection, and 2-, 4- and 7-DPI to assess virus shedding. The swabs were placed into 2 ml of virus transport medium comprising tissue culture medium 199 (Sigma-Aldrich) supplemented with 25 mM 4-(2-hydroxyethyl)-1 piperazineethanesulfonic acid (HEPES), 0.035% sodium bicarbonate, 0.5% bovine serum albumin, penicillin 100 IU/ml, streptomycin 100 μg/ml, and nystatin 0.25 μg/ml, vortexed and centrifuged to remove debris, then stored at −80°C for subsequent virus titration. Blood samples were collected in BD Vacutainer SST™ serum separation tubes at the start of the study (prior to challenge) and at 14-DPI. BAL was collected from the entire left lung with 0.1% BSA in PBS. BAL samples were centrifuged at 500 × g for 5 min and the supernatant was aliquoted and frozen for further analysis.

### Plaque assay

Virus titres in nasal swabs were determined by plaque assay on MDCK cells as described previously [22]. Plaques were visualized by staining the monolayer with 0.1% (v/v) crystal violet containing 20% methanol (Merck) and expressed as plaque forming units (PFU) per ml of nasal swab eluate.

### Virus whole-genome sequencing

Viral RNA extraction and influenza virus whole genome sequencing was performed as previously described [23]. Raw fastq sequences were then imported into Geneious Prime software and, trimmed to remove primers and aligned to whole genome reference sequences (GM19 [EPI_ISL_377080], Syd21 [EPI_ISL_19193801], Wsc19 [EPI_ISL_404527], Lis23 [EPI_ISL_18788764], Wsc22 [EPI_ISL_15928538], Vic22 [EPI_ISL_19192400]).

### Ferret antisera production

Post-infection ferret antisera were generated by intranasally inoculating ferrets with 500 μl (250 μl per nostril) of virus. Two weeks after inoculation sera were collected.

### Human serology panels

Pre- and two-week post-vaccination human sera were collected over three consecutive years from healthy volunteers working at the Francis Crick Institute. Inclusion criteria included the receipt of a seasonal vaccine, being employed at the Francis Crick Institute and consenting to give a pre- and post-vaccination blood sample. No details of previous vaccination/infection history were obtained. Samples were collected in BD Vacutainer SSTII tubes and centrifuged for 10 minutes at 2000rpm to separate red and white blood cells from the serum. The serum supernatants were transferred into separate tubes and heat inactivated at 56°C for 30 minutes before storage at −20°C.

### Hemagglutination Inhibition assay (HI)

Ferret and pig antisera were heat inactivated at 56°C for 30 minutes and treated with receptor destroying enzyme (RDE) (Denka Co.). RDE-treated sera were diluted to a working dilution of 1:20 in PBS with 0.1% azide.

Hemagglutination and hemagglutinin inhibition (HI) assays were performed according to standard methods using suspensions of turkey RBCs (0.75% v/v) [24]. Four hemagglutination units were used in all HI assays.

### Virus neutralisation (VN)

Viruses were titrated to determine the dilution at which 50% of cells within a well of a 96-well plate were infected, these dilutions were used in the VN assay as previously described [25].

## Author contribution

ET, IB, NL, RH, RD, JM designed the study and obtained the funding; ET, IB, TC, BP, CH, EB designed and performed the pig animal experiments; TC, BP, CH, EB, SR, processed samples; RH, TC, BP, AL, CC, TP, acquired, analysed and interpreted the data; ET, IB, JM, RH, RD wrote the manuscript. All authors have read and approved the manuscript.

## Acknowledgments

We are grateful to the animal staff at the APHA for providing excellent animal care. This work was supported by the UKRI Biotechnology and Biological Sciences Research Council (BBSRC) IAA award BB/X511134/1 Research at Pirbright was funded by BBSRC via the Pirbright Institute’s Strategic Programme Grants (ISPGs) [BBS/E/PI/230002A; BBS/E/PI/230002B], BBSRC National Bioscience Research Infrastructure: High Containment and Low Containment Services and Science Platforms grants [BBS/E/PI/23NB0004, BBS/E/PI/23NB0003]. This work was supported by core funding from the Francis Crick Institute from Cancer Research UK (grant no. CC1114), the UK Medical Research Council (grant no. CC1114), and the Wellcome Trust (grant no. CC1114). SR, TP and IB are supported by the Biotechnology and Biological Sciences Research Council (BBSRC)/DEFRA ‘FluTrailMap’ consortium [BB/Y007298/1] and the UK Medical Research Council/Department for Environment, Food and Rural Affairs (Defra, UK) FluTrailMap-One Health consortium [MR/Y03368X/1]. TC was supported by Gates Foundation grant OPP1215550 (Pirbright Livestock Antibody Hub).

## Ethics statement

The pig studies were approved by the ethical review processes at APHA and the Pirbright Institute in accordance with the UK Government Animal (Scientific Procedures) Act 1986 under Project License PP2064443. Collection of human pre- and post-vaccination serum samples was approved by The Francis Crick Institute Internal Ethics Review Committee, registration number: 2019 FC5.

